# Viral infection to the raphidophycean alga *Heterosigma akashiwo* affects both intracellular organic matter composition and dynamics of a coastal prokaryotic community

**DOI:** 10.1101/2024.10.07.616994

**Authors:** Hiroaki Takebe, Haruna Hiromoto, Tetsuhiro Watanabe, Keigo Yamamoto, Keizo Nagasaki, Ryoma Kamikawa, Takashi Yoshida

## Abstract

Marine microalgae play a crucial role in marine ecosystem by supplying dissolved organic matter to heterotrophic prokaryotes, which mediate the microbial loop. Microalgae are often infected by viruses, and in infected cells (virocells), the viruses modulate and often change the host metabolism for propagation and thereby endo-metabolites. However, the impact of algal virocells on prokaryotic communities is not fully been understood. In this study, we investigated whether lysates from virocell of *Heterosigma akashiwo,* a globally prevalent bloom-forming raphidophycean alga, causes shifts inn prokaryotic community structure, and which metabolic compounds in the viral lysate might affect the surrounding prokaryotic populations. Using microcosm experiments, we cultured prokaryotic communities with a dissolved fraction derived from the viral lysate (VDF) of *H. akashiwo*. Results revealed that certain prokaryotic populations assigned as the *Vibrio* spp. pathogenic against fish and crustaceans grew specifically responding to the VDF. These *Vibrio* species possessed a gene module for branched-chain amino acids transporters, which were revealed to be enriched in VDF by gas chromatography-mass spectrometry analysis. Altogether, our findings suggest that viral infection-induced changes in biochemical properties of *H. akashiwo* cells can promote the growth of taxonomically and metabolically different prokaryotic populations, potentially impacting higher trophic-level consumers in marine ecosystems.

**Highlights:**

- The effect of viral infection to microalgae on marine prokaryotes was investigated.
- Viral lysate of *Heterosigma akashiwo* promoted growth of certain *Vibrio* population.
- Those *Vibrio* likely have ability to utilize the compounds enriched in the lysate.
- Those *Vibrio* have been known as pathogen of fish or crustacean.
- Viral infection to *H. akashiwo* can indirectly affect higher trophic-level consumers.

## 1. Introduction

Marine microalgae significantly contribute to the primary production. Of the produced organic matters, the dissolved fractions (dissolved organic matter: DOM) are mainly consumed by planktonic heterotrophic prokaryotes (Azam et al., 1983; Field et al., 1998; Moran et al., 2016). Viruses that infect these algae are also ecologically important, as they play contribute to the mortality of host cells and the release of organic matter from the cells, thereby affecting biogeochemical cycles (Suttle, 2005, 2007). Additionally, while particles of viruses floating in the seawater are regarded as those under a metabolically passive state, virus-infected host cells have been considered as a “living forms” of the viruses and thus referred to as virocells (Forterre, 2013). Viruses infecting host cells actively modulate host cell metabolisms through production of their daughter virions (Forterre, 2013; Zimmerman *et al*., 2020).

The nucleo-cytoplasmic large DNA virus (NCLDV), a group of eukaryotic viruses, builds unique cytoplasmic structures, such as the viral factory and autophagosome, in their virocells. These structures lead to the production of unique endo-metabolites that differ from those in uninfected cells (Forterre, 2013; Rosenwasser *et al*., 2016). For example, in virocells of the marine, bloom-forming haptophyte *Emiliania huxleyi,* the production of a specific sphingolipids that are integral to viral particles is highly upregulated (Rosenwasser *et al*., 2014). Similar virocell-specific metabolic changes have been reported in the harmful bloom-forming pelagophyte *Aurerococcus anophagefferens* and the globally distributed pico-sized chlorophyte *Ostreococcus lucimarinus* (Moniruzzaman *et al*., 2018; Zimmerman *et al*., 2019). Given that the preferred organic matter of heterotrophic prokaryotes varies among species (Alonso-Sáez and Gasol, 2007; Sarmento *et al*., 2013), virocells are likely to have different effects on prokaryotic community dynamics compared to the uninfected cells. Indeed, viral infection of *E. huxleyi* have been shown to cause significant shifts in natural prokaryotic communities in the mesocosm bags in coastal areas, though the specific virocell-derived organic matters were not analyzed inn these studies (Vincent *et al*., 2023).

The raphidophycean alga *Heterosigma akashiwo* represents a crucial model for investigation of ecological relationships among microalgae, their viruses, and the surrounding prokaryotes. *H. akashiwo* is known for causing harmful blooms that result in mass mortality of fish and shellfish globally (Chang *et al*., 1990; Nagasaki *et al*., 1994; Needham and Fuhrman, 2016; Matcher *et al*., 2021). Our previous study demonstrated that particular prokaryotic populations, such as those from *Alteromonadales* and *Vibrionales,* respond to DOM extracted from uninfected *H. akashiwo* cells in microcosm experiments (Takebe *et al*., 2020). These responses differed from those observed with DOM from viral-free algal cells of a diatom, suggesting that blooms of different primary producers result in distinct prokaryotic community dynamics (Takebe *et al*., 2024). Importantly, *H. akashiwo*-infecting viruses (HaVs) have been isolated during natural *H. akashiwo* blooms (Nagasaki and Yamaguchi, 1998; Nagasaki *et al*., 1999; Tarutani *et al*., 2000). Notably, up to 10^5^ cells of *H. akashiwo* per mL of surface seawater can disappear within 3 days, accompanied by an increase in extracellular HaV particles (Tarutani *et al*., 2000). If HaVs are the primary drivers behind the decline of *H. akashiwo* blooms, substantial amounts of organic compounds are expected to be released from *H. akashiwo* virocells during the final stages of the bloom. However, the impact of *H. akashiwo* virocells on surrounding prokaryotic community structures remain to be investigated.

In this study, we investigated how organic compounds derived from *H. akashiwo* virocells affect prokaryotic community dynamics. Organic compounds released from microalgal cells can be categorized into three different modes, exudates from healthy living cells, intracellular components released due to grazing, and viral-mediated cell lysates (Ma *et al*., 2018). To investigate these effects, we conducted microcosm experiments using a prokaryotic community collected from surface water in Osaka Bay. The experiments involved adding viral lysates of *H. akashiwo*, exudates, or intracellular components from uninfected *H. akashiwo*. We employed 16S rRNA gene amplicon analysis to assess prokaryotic community shifts and Gas Chromatography-Mass Spectrometry (GC-MS) to analyze DOM derived from both virocell and uninfected cells. Our findings revealed that virocell lysates, which had a unique organic matter composition, led to an increase in specific bacterial populations that appear to utilize these organic matters as substrates. Our data suggest that viral infection induced-changes in organic compounds may promote growth of prokaryotes with specialized metabolic capacities.

## 2. Materials and Methods

### 2.1. Culturing of *H. akashiwo*

*H. akashiwo* strain NIES-293 was purchased from the National Institute for Environmental Studies (NIES) and was maintained axenically in f/2 medium (Guillard and Ryther, 1962) at 20 °C with a 12-hour light/12-hour dark photoperiod at an irradiance of 40 µmol photons m^-2^ s^-1^. The *culture* was grown in 1.5L of medium under these conditions. It was then divided into three portions (500 mL, 600 mL, and 400 mL), for the following purposes: extraction of exudates and intracellular components from uninfected cells, preparation of viral lysates, and monitoring of growth until the stationary phase, respectively.

### 2.2. Collection of exudates and intracellular components

To prepare intracellular components and exudates from uninfected cells, 500 mL of the culture was centrifuged at 1,500 x g for 5 min using a High Capacity Bench-top Centrifuge LC-220 (TOMY SEIKO). The resulting supernatant was filtered through a 0.2-µm pore size PVDF filter (Millipore), and the filtrate was designated as the exudate-derived dissolved fraction (EDF). Intracellular components was extracted from the pellet using the method outlined in our previous study (Takebe *et al*., 2024) yielding the intracellular components-derived dissolved fraction (IDF).

### 2.3. Viral infection experiments of *H. akashiwo*

The *H. akashiwo*-infectious virus strain HaV103, isolated from Uranouchi Bay, Kochi, Japan in 2021, was inoculated in 600 ml of the NIES-293 culture with an initial multiplicity of infection (MOI) of approximately 0.008, to prepare viral lysates. Sixty mL of the infected culture was reserved for monitoring the dynamics of NIES-293 and HaV103. After the culture was decolorized, viral lysates were harvested from the remaining 540 mL . The supernatant was filtered through a 0.2-μm PVDF filter (Millipore). The resulting filtrate was designated as the viral lysates-derived dissolved fraction (VDF). The VDF was then stored at −80°C until further use.

### 2.4. Cell counting of *H. akashiwo* and enumeration of extracellular HaV

To count *H. akashiwo* cells, 1,920 µL of culture medium was analyzed using RF-500 Flow Cytometer (Sysmex Corporation). Cells was quantified by plotting chlorophyll fluorescence in the red channel (695/50 nm) against forward scatter, and the data were analyzed with FlowJo (Becton, Dickinson and Company) following the manufacturers’ instructions.

For enumerating extracellular HaV particles, 20 mL of the infected *H. akashiwo* culture was transferred each of three flasks and then diluted 10-fold with fresh f/2 medium. From each diluted culture, 12 mL was centrifuged at 1,500 x g for 5 min to remove host cells and debris. The supernatant was then centrifuged again under the same conditions. The final supernatant was ultracentrifuged at 104,000 × *g* using an Optima XE-90 Ultracentrifuge (Beckman Coulter Inc.) to pellet the viral particles. Viral DNA was extracted from the pellet using the xanthogenate-SDS method described by Kimura *et al*. ( 2012). The abundance of HaV103 was assessed by quantifying the gene encoding the major capsid protein (mcp) using quantitative PCR. Primers specific for the mcp gene (HaV53_mcp_290F: 5′-GTCTGAGGACGCACTGTGAA and HaV53_mcp_429R: 5′-GCTGGTACTGCGTACAACGA) were designed from the genome sequence of HaV53 (the GenBank accession number KX008963; Ogura *et al*., 2016). These primers were tested for specific amplification of the target gene using DNA from *H. akashiwo* NIES-293 and HaV103 analysis. PCR reactions were conducted with TB Green® Premix Ex Taq™ II (TaKaRa Bio) as per the manufacturer’s instruction. The Thermal Cycler Dice® Real Time System III (TaKaRa Bio) was used with the following cycling conditions: 1 cycle at 95 for 30 s, followed by 40 cycles of 95 for 5 s, 60 for 30 s, and 72 for 15 s. A standard curve for quantifying viral particle abundance was constructed using serial dilutions of known concentrations of HaV103 DNA .

### 2.5. Measurement of carbon concentration in EDF, IDF, and VDF

To measure the carbon concentration in the VDF, IDF, and EDF, as well as f/2 medium (used as a control), approximately 20 ml of each sample was analyzed using Non-Purgeable Organic Carbon analysis with a Total Organic Carbon Analyzer TOC-L, Shimadzu).

### 2.6. Setup of microcosm experiments

On June 22, 2022, approximately 10 L of surface seawater (5 m depth) was collected from Osaka Bay, Japan (N 34°19’28”, E 135°7’15”). The seawater was pre-filtered through polycarbonate membrane filters (142 mm diameter, 3.0-μm pore size; Millipore) to remove eukaryotic cells. For the microcosm experiments, 4.8 L of the pre-filtered seawater was again filtered through polycarbonate membrane filters (142-mm diameter, 0.2-μm pore size; Millipore) to isolate coastal prokaryotic cells. The filters capturing the prokaryotic fraction was divided into four equal pieces, and the cells on each piece were re-suspended in 300 mL of autoclaved aged seawater in 500 mL flasks. The residual organic matter in the flasks was removed by washing with 6 N HCl followed by Milli-Q water.

VDF was added to three flasks containing the prokaryotic fraction to a final concentration of 80 µmol C/L, which is representative of carbon concentrations observed in natural microalgal blooms (Børsheim *et al*., 1999). IDF, EDF, and f/2 medium (used as a control) were applied in the same manner as VDF. The flasks were categorized into four treatment groups: control, VDF-treatment, IDF-treatment, and EDF-treatment. Each treatment consisted of three triplicate flasks, designated as replicate flasks I-III. A total of twelve flasks were incubated at 20°C, under the 14.5-hour light/9.5-hour dark cycle at an irradiance of 150 µmol photons m^-2^ s^-1^, for 4 days. These temperature and light conditions represent the typical environmental conditions of bloom-forming seasons at Osaka Bay, Japan.

Control, IDF-treatment, EDF-treatment, and VDF-treatment flasks were homogenized by mixing once daily before subsampling. Each day, 1,920 µl was subsampled from each flask every day and fixed with glutaraldehyde (1% final concentration) at 4°C. The fixed prokaryotic cells were stained with SYBR® Green I (final concentration 1×; Thermo Fisher Science) for 20 minutes at room temperature in the dark. The stained cells were then counted using an S3e Cell Sorter (Bio-Rad) and analyzed using FlowJo (Becton, Dickinson and Company), according to manufacturers’ instructions. Mann–Whitney U tests were performed to compare cell and viral abundances among treatments on the same day .

### 2.7. DNA extraction and sequencing

For prokaryotic community structure analysis, 1,800 µl of culture medium was subsampled daily from each flask, resulting in a total of 60 samples over 4 days. Prokaryotic cells were collected on polycarbonate membrane filters (25-mm diameter, 0.2-μm pore size; ADVANTEC Toyo Kaisha, Ltd.) and stored at −30°C until DNA extraction. To analyze the prokaryotic community composition in the original seawater, 10 ml of the seawater was filtered on polycarbonate filters (25-mm diameter, 0.2-μm pore size; ADVANTEC Toyo Kaisha, Ltd.), and these filters were stored at −30 °C until DNA extraction. DNA was extracted using the method described in our previous study (Takebe *et al*., 2020). The control flask replicate I sample on day 2 was lost due to a technical error. The 16S rRNA genes were amplified using primers targeting the V3–V4 hypervariable regions, with added overhang adapter sequences at each 5 -end, as described by (Takahashi *et al*., 2014) and according to the Illumina 16S sample preparation guide. The amplicons were sequenced with a MiSeq Reagent kit, version 3 (2×300 bp read length; Illumina), following to the manufacturer’s instruction.

### 2.8. Sequence processing and ASV generation

From the original seawater sample, 208,405 reads were obtained (Supplementary Table 1). During the microcosm experiments, the average number of reads per flask were as follows: 1,019,575 for the control, 1,026,863 for the VDF-treatment, 1,048,035 for the IDF-treatment, and 952,122 for the EDF-treatment (Supplementary Table 1). Quality filtering, denoising, pair-end merging, and construction of the amplicon sequence variants (ASVs) feature table were performed using the DADA2 plugin (Callahan *et al*., 2016) within QIIME2. Taxonomic assignment of the ASVs was performed using the SILVA (release 138) reference database (Quast *et al*., 2013). ASVs assigned to mitochondrial or chloroplast sequences were removed from the feature table and excluded from subsequent analyses. Singleton ASVs were also removedduring this process.

### 2.9. Analysis of community structure and dynamics of prokaryote ASVs

To analyze ASV composition, all sequence data were rarefied to the depth of the lowest sample. The Bray-Curtis dissimilarity index was calculated for pairwise comparisons of the prokaryotic communities using the “vegan” package in R. This index was visualized by principal coordinate analysis (PCoA) with the “stats” package in R. The Analysis of similarity (ANOSIM) was performed with the Bray-Curtis dissimilarity scores to assess significance of differences among days or treatments, using the “vegan” package in R. *P*-values for multiple comparisons were corrected using the Bonferroni method.

Relative abundance of each ASV was calculated by dividing the read counts of the ASV by the total read counts of each sample. To estimate the approximate cell density (cells/mL) of each ASV, the relative abundance (ranging from 0 to 1) was multiplied by the total number of prokaryotic cells. ASVs were classified as abundant if they ranked among the top 20 in approximate cell density at least one day after day 1 in all the triplicate flasks, and if their maximum approximate cell density was more than twice that at day 0. Differences in the approximate cell density of these ASVs among treatments were tested for statistical significance using LEfSe (Segata *et al*., 2011), with treatment as the class and triplicate flasks as subclass.

### 2.10. Survey of abundant Osaka Bay ASVs in amplicon dataset from Monterey Bay natural bloom samples

To confirm that the ASVs of interest were detected in the natural *H. akashiwo* bloom, we analyzed abundance of these ASVs and *H. akashiwo* during a bloom observed in Monterey Bay, US. The 18S rRNA gene sequence of *H. akashiwo* NIES-293 (DQ470658.1) (Ki and Han, 2005) was obtained from GenBank (Benson *et al*., 2012). Raw amplicon reads of 18S rRNA and 16S rRNA gene sequences from natural bloom samples collected daily in Monterey Bay between September 26 and November 16, 2016 (Nowinski *et al*., 2019) were retrieved from the NCBI Sequence Read Archive (PRJNA533622). These reads were quality-trimmed, merged, and chimeras were removed following the pipeline described in our previous study (Takebe *et al*., 2020). Samples with fewer than 5,000 quality-controlled reads were excluded from the analysis. The quality-controlled 18S rRNA and 16S rRNA reads were mapped to the NIES-293 sequence and prokaryotic ASVs of interest with 97% and 100% identity, respectively, using VSEARCH (Rognes *et al*., 2016). Relative abundance was then calculated using the method described above. Sequences with 97% or higher identity to the 18S rRNA gene of NIES-293 were considered to be *H. akashiwo*.

### 2.11. Phylogenetic analysis of abundant ASVs

To determine the phylogenetic positions of the abundant ASVs of interest, nearly complete 16S rRNA gene sequences (> 1500 bp) of their assigned genera, as identified by Silva (see above), were retrieved from the SILVA NR reference database to construct reference phylogenetic trees. Sequences were aligned using MAFFT (Katoh and Standley, 2013) with the L-INS-i method, and gap positions were automatically removed using trimAl (Capella-Gutierrez *et al*., 2009) using the -gappyout option. Phylogenetic tree were then reconstructed using the approximately maximum-likelihood method using FastTree (Price *et al*., 2010). The abundant ASVs of interest were classified onto the reference tree using pplacer (Matsen *et al*., 2010). Results were visualized with iTOL (Letunic and Bork, 2019).

### 2.12. Comparative genomic analysis of abundant ASV-relatives

Complete genomes of the closest isolates (100% identity of 16S rRNA sequence) of the ASVs of interest were downloaded from the NCBI RefSeq NR database (Pruitt *et al*., 2007). KEGG modules present in these genomes were identified using METABOLIC (Zhou *et al*., 2022) with default settings. The presence or absence of each module in each genome was used to calculate the possession rate among ASV-relative lineages.

### 2.13. Metabolomic analysis using gas chromatography-mass spectrometry (GC-MS)

An internal control (2-isopropylmalate) was added to the 250 µL each of sodium chloride (26.4 mg mL−1) as the blank, f/2 medium (control), VDF, IDF, and EDF. The samples were freeze-dried for over 6 h. To remove residual water, 250 μL of toluene (> 99.7%) was added to each sample, and the mixture was ultrasonicated for 10 min. The toluene was removed under a gentle flow of N2 gas. Metabolite derivatization was performed by adding 80 μL of methoxyamine hydrochloride (98%) dissolved in pyridine (20 mg/mL) to the dried pellet. The mixture was ultrasonicated for 10 min, briefly vortexed to dissolve the pellet, and incubated for 90 min at 30°C with constant rotation at 1,200 rpm in a thermal rotating incubator. Following this, 100 μL of trimethylsilyl-N-methyl trifluoroacetamide was added, and the mixture was ultrasonicated for 10 min, vortexed, and incubated for 30 min at 37°C with constant rotation at 1,2000 rpm. The derivatized mixture was ultrasonicated for an additional 10 min, and residual salts were removed by centrifugation at 16,000 × g for 5 min at room temperature. The supernatant was then analyzed by GC-MS.

All derivatized samples were analyzed on GCMS-QP2010Ultra (Shimazu). Retention times were calibrated with the Qualitative Retention Time Index Standard (AART-STD; Restek Corporation) prior to analysis. The samples were injected onto a DB-5 column (30 cm by 0.25 mm, 1.0-μm film thickness; Agilent). The injector temperature was set at 280°C. Chromatographic separation was achieved with an initial column oven temperature of 100°C (held for four minutes), followed by a temperature ramp of 10°C/min until reaching 320°C, which was then maintained for 11 min. Helium was used as a carrier gas at a constant flow rate of 39 cm/sec. Mass spectra were collected in electron ionization mode at 70 V across the mass range of m/z 45 to 600.

For peak annotation in the GC-MS analysis, 401 compounds from the SHIMADZU Smart Metabolites Database (Shimadzu) were used as references. The relative abundance of each detected compound was calculated by dividing its peak area by that of the internal control. This relative abundance was then normalized to account for the carbon concentration in each sample, allowing for accurate comparison between samples with different carbon concentrations.

## 3 Result

### 3.1. Abundance shifts of *H. akashiwo* and HaV during infection experiments

Since microalgal cell death occurs upon viruses infection without the concomitant release of daughter virions (Vincent *et al*., 2021), it is substantial to confirm that cell lysis is due to viral infection to obtain genuine viral lysates. In our study, cultures of *H. akashiwo* NIES-293 infected with HaV103, including those at a 10-fold dilution, exhibited growth arrest by day 3 (1.63 ± 0.18 × 10^5^ and 7.63 ± 1.16 × 10^4^ cells/mL, respectively) and subsequently lysed by day 6. In contrast, uninfected cultures continued to grow until day 14 (Supplementary Fig. 1A). Extracellular viral DNA accumulation began as early as day 3 in the 10-fold diluted infected culture and peaked on day 5 (1.54 ± 1.22 × 10^7^ *mcp* copies/mL) (Supplementary Fig. 1B), indicating the release of HaV particles. Therefore, the observed lysis of *H. akashiwo* NIES-293 was attributable to viral infection. These observations indicated that the resultant lysate was derived from genuine HaV-infected *H. akashiwo* virocells, which were used for subsequent experiments.

### 3.2. Abundance shifts of prokaryotic cells in microcosms

We investigated the effects of dissolved fraction of viral lysate of *H. akashiwo* NIES-293 (VDF) on prokaryote cell numbers in the natural seawater from Osaka Bay. Cell densities across all treatments showed similar dynamics and abundances (Fig. 1). All treatments showed a sharp increase in cell densities from day 1 to day 3, reaching maximum densities on day 4 (2.56 ± 0.36 × 10^6^ – 4.08 ± 0.61 × 10^6^ cells/ml) followed by stagnation (Fig. 1). The cell densities in the VDF-treatment did not significantly differ from those in the EDF-treatment and IDF-treatment on day 4, indicating that the effect of the *H. akashiwo* NIES-293 virocell lysate on prokaryotic abundance was comparable to that of intracellular components or exudates from uninfected *H. akashiwo* NIES-293 cells. These results indicate that certain prokaryotes grew in these conditions.

**Fig. 1.**
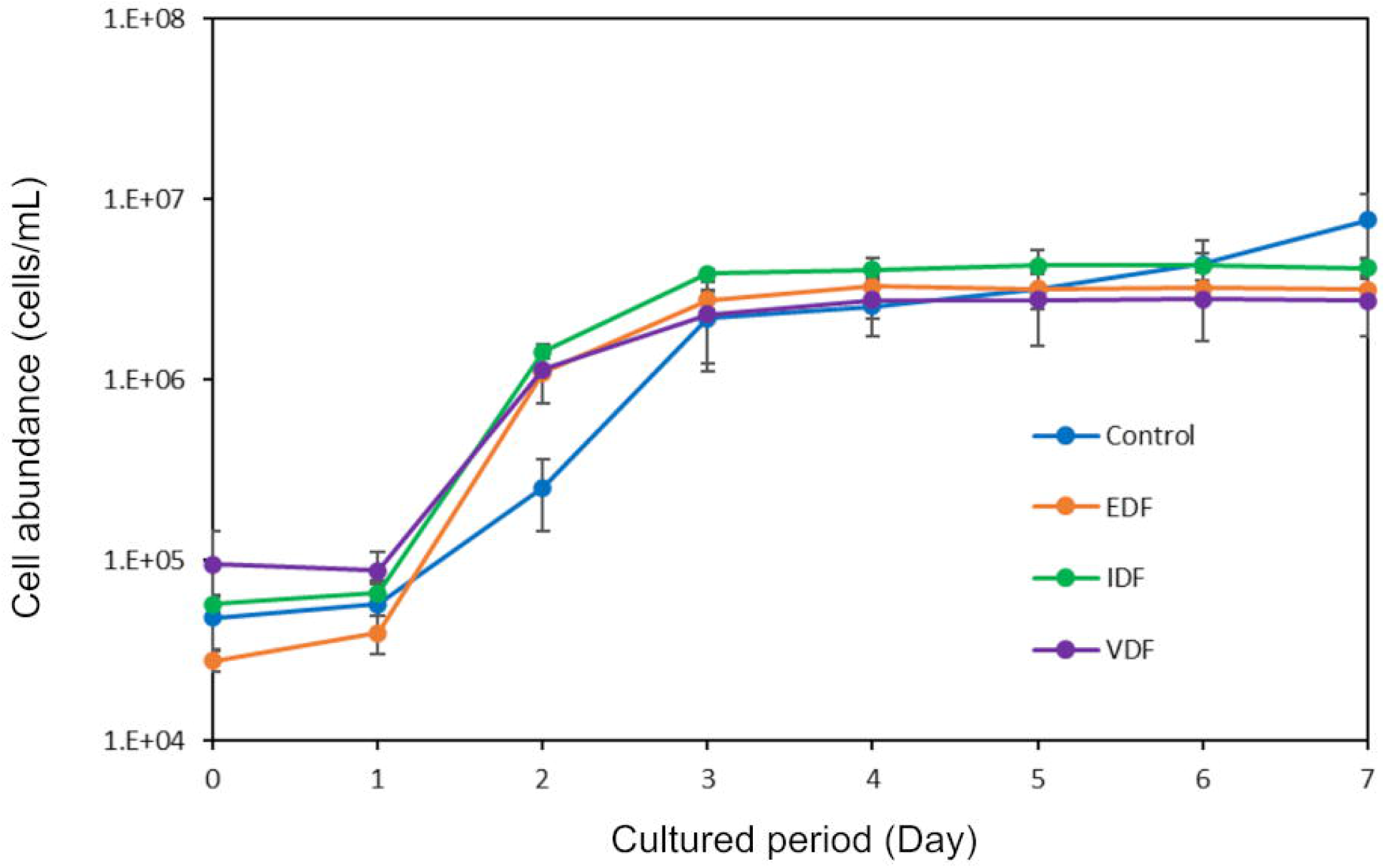
Shifts in abundance of prokaryotic cells during the microcosm experiment. Cell counts were obtained using flow cytometry. The average cell and viral numbers across triplicate flasks are shown. Error bars indicate standard deviation.

### 3.3. Shifts in prokaryotic community structure in response to VDF

Next, we investigated the effect of VDF on the marine prokaryotic community structure using 16S rRNA gene amplicon sequencing. From the original seawater sample, we identified 606 ASVs. During the microcosm experiments, the average number of ASVs obtained per flask was 1,080 for the control, 893 for the VDF treatment, 385 for the IDF treatment, and 976 for the EDF treatment (Supplementary Table 1). We examined whether the effect of VDF on ASV compositions differed from that of IDF and EDF using principal coordinate analysis (PCoA) based on Bray-Curtis dissimilarity (Fig. 2). While day 0 samples were closely clustered with the original seawater sample, samples from day 2 to day 4 in the same treatments were more similar to each other but distinct from those in other treatments (ANOSIM, *p* < 0.05) and from day 0 samples within the same treatments (ANOSIM, *p* < 0.05) (Fig. 2). This PCoA indicates that prokaryotic community structures shifted differentially depending on the treatment and cultivation time. Especially, VDF appeared to influence prokaryotic community structures, creating a distinct profile compared to IDF and EDF.

**Fig. 2.**
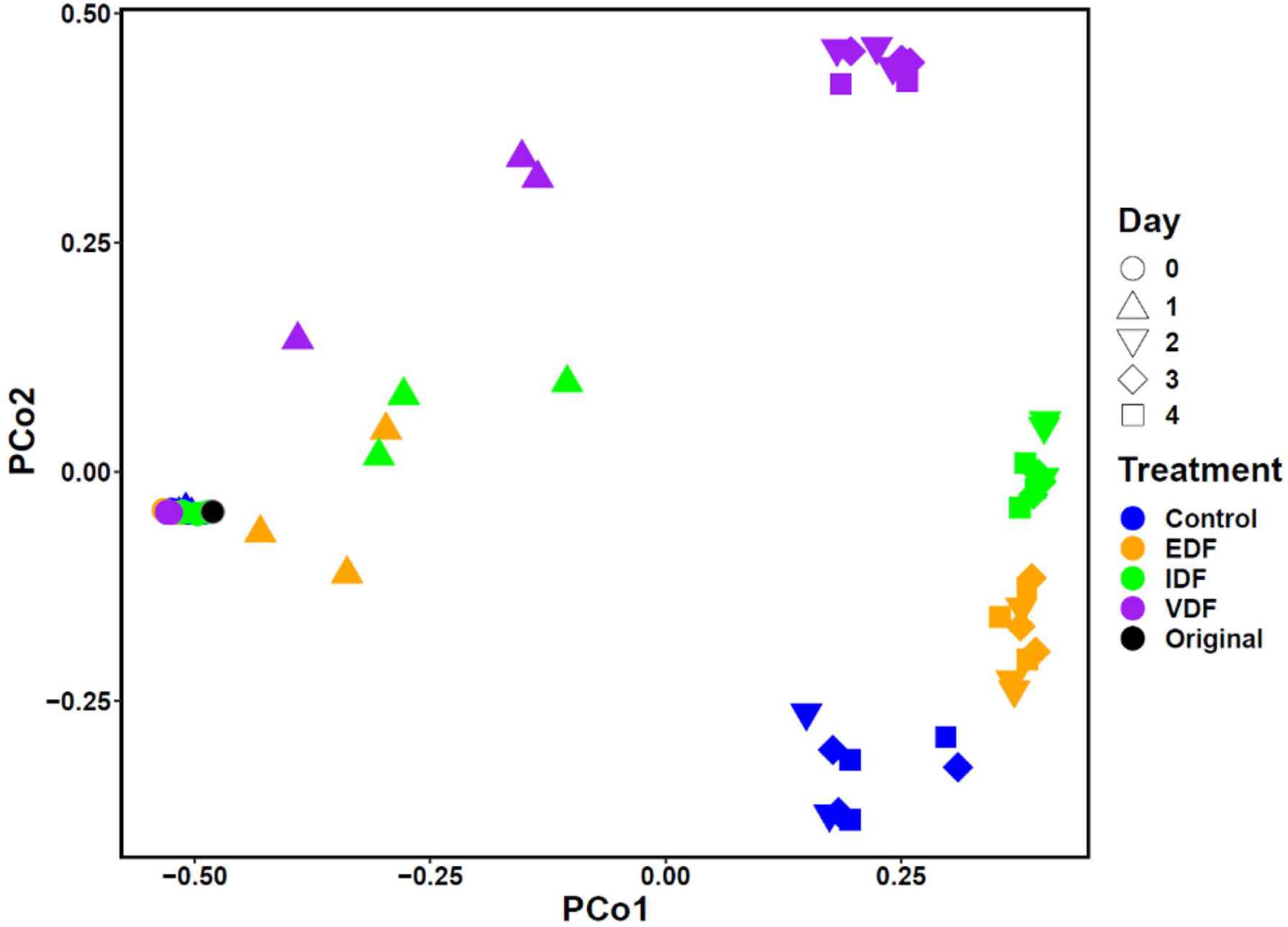
Comparison of ASV compositions across control, VDF-, IDF-, and EDF-treatments. The number of sequences of each sample were rarified into 3,453 reads prior to the analyses. Bray–Curtis dissimilarity among all samples was visualized using Principal Coordinate Analysis (PCoA). Samples are distinguished by colors and shape according to their treatments and culture periods, respectively.

We further analyzed which phyla (class level for Proteobacteria) became dominant in the VDF treatment (Supplementary Fig. 2). On day 0, the dominant heterotrophic prokaryotes in the VDF samples of were Alphaproteobacteria and Gammaproteobacteria (24.5% and 24.7%, respectively), followed by Bacteroidetes (15.2%), whereas smaller proportions were of Actinobacteria, Verrucomicrobia, and Thermoplasmatota. Alphaproteobacteria, Gammaproteobacteria, and Bacteroidetes remained abundant throughout the microcosm experiment, their relative abundances shifted during the experiment. By day 2, Gammaproteobacteria dominated almost entirely, while the proportions of Alphaproteobacteria and Bacteroidetes decreased. The increase in the total prokaryotic cell count in the VDF-treatment (Fig. 1) suggest that lysates from *H. akashiwo* NIES-293 virocells specifically promoted the growth of certain populations in Gammaproteobacteria populations (Supplementary Fig. 2). Similar shifts in community structure were also observed in EDF and IDF treatments, although the proportions of dominant phyla or classes varied (Supplementary Fig. 2). In contrast, Campylobacterota became more dominant in the control treatment with added f/2 was added. These shifts at the phylum- and class-level were more explicit from day 2 and consistent with the PCoA patterns (Fig. 2).

### 3.4. Abundant ASVs in VDF and the other treatments

Although the PCoA analyses indicate distinct community structure shifts among VDF, IDF, and EDV, treatments, the phylum- or class-level proportions were not largely distinct across the treatments (Supplementary Fig. 2). This suggests that the differences in community structure would be observed at finer taxonomic level. Therefore, we focused on prokaryotes that specifically responded to and grew on VDF at the ASV level. Since changes in relative abundance during cultivation do not necessarily reflect cell growth and death, we analyzed the dynamics of abundant ASVs by calculating their approximate cell density, which combines the total cell number and relative abundance.

We identified 11, 16, 17, and 13 abundant ASVs in the control, IDF-, EDF-, and VDF-treatments, respectively (Supplementary Fig. 3). Of the 13 ASVs abundant in the VDF-treatment (Fig. 3), 7 were specifically abundant in the VDF-treatment. These included three ASVs from the Vibrionaceae family (ASV_3626; genus not assigned, ASV_3968; *Vibrio*, and ASV_4145; *Vibrio*), three ASVs from the genus *Pseudoalteromonas* (ASV_1731, ASV_4440, and ASV_927), and one ASV from *Psychrobirium* (ASV_1418) (Fig. 3; Supplementary Table 2). All these ASVs belong to Gammaproteobacteria. Differential abundance testing using Lefse indicated that all these ASVs, except for *Pseudoalteromonas* ASV_927, were significantly more abundant in the VDF-treatment compared to the other treatments (Supplementary Table 2). Hereafter, the 6 ASVs significantly more abundant in the VDF-treatment were referred to as VDF-specific ASVs.

**Fig. 3.**
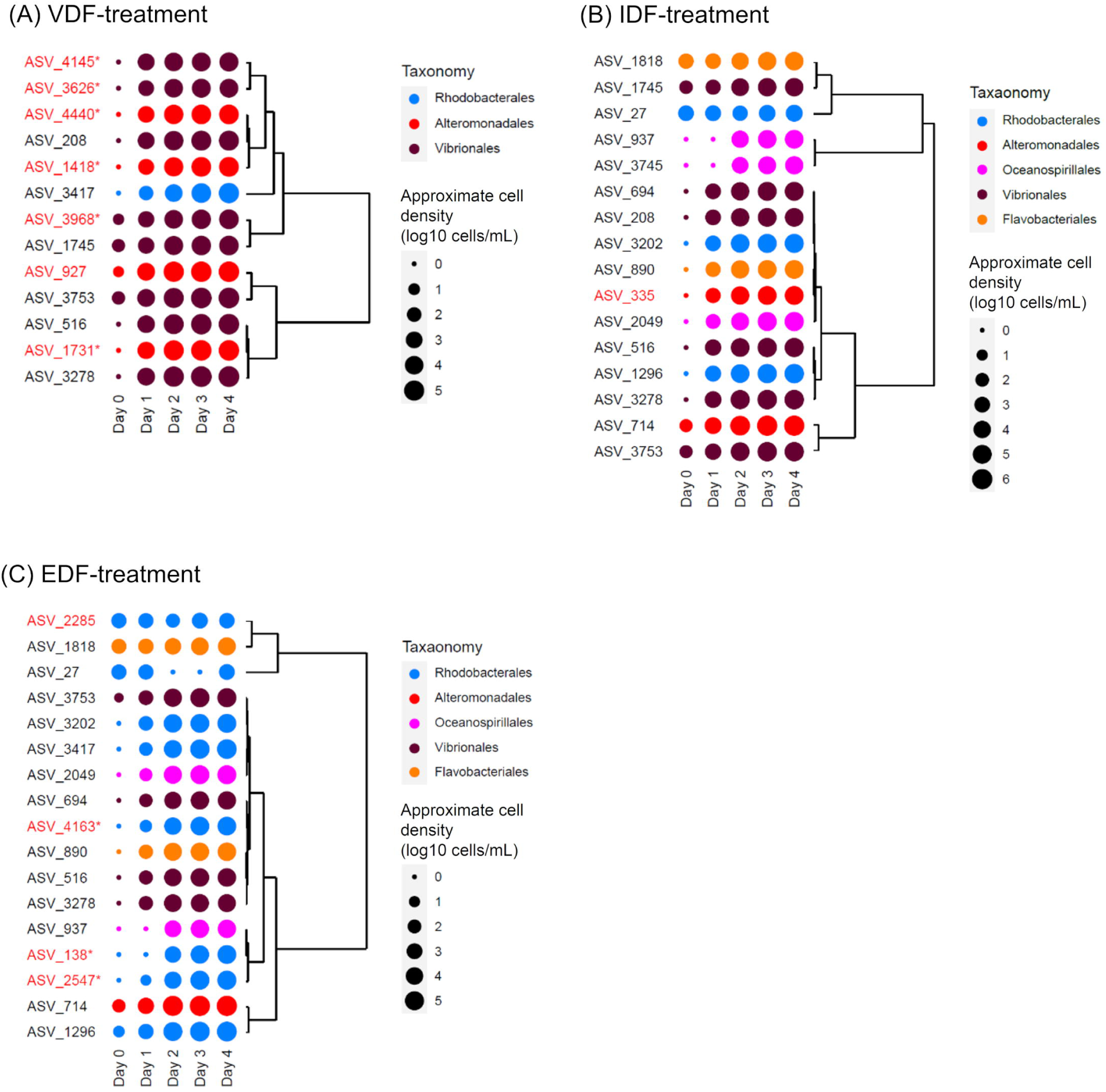
Dynamics of ASVs abundant in (A) VDF-treatment, (B) IDF-treatment, and (C) EDF-treatment. The plots show average approximate cell density in the triplicate flasks on a log scale. Colors show order-level taxonomy of each ASV. Dendrograms represent similarity in approximate cell density dynamics among ASVs. ASVs abundant exclusively in VDF-, IDF-, or EDF-treatments are highlighted in red, with treatment-specific ASVs indicated by asterisks.

The remaining 6 abundant ASVs in the VDF treatment were also found to be abundant in at least one of the other treatments (Supplementary Fig. 3; Supplementary Table 2). These shared ASVs were assigned to the genera *Thalassibius* (ASV_3417) and *Vibrio* (the other ASVs) based on the Silva database (Supplementary Table 2). Additionally, 9 ASVs abundant in either or both of IDF and EDF were considered capable of utilizing dissolved fractions from uninfected *H. akashiwo* cells. Among these, *Pseudoalteomonas* ASV (ASV_335) and 4 of Rhodobacteraceae ASVs (ASV_138; genus not assigned, ASV_2285; *Planktomarina*, ASV_2547; genus not assigned, and ASV_4163; Octadecabacter) were abundant exclusively in the IDF- and EDF-treatment, respectively, (Supplementary Table 2). The comparison of abundant ASVs highlighted the distinct effects of dissolved fractions from virocells and uninfected *H. akashiwo* cells on prokaryotic community structures at the ASV level, even within the same genera. Hereafter, ASV_927, shared ASVs, and ASVs abundant in the treatments other than VDF will be referred to as VDF-nonspecific ASVs.

### 3.5. Taxonomic and phylogenetic relationships between VDF-specific and -nonspecific ASVs

As most of VDF-specific and -nonspecific abundant Vibrionaceae ASVs (3 and 6 ASVs, respectively) were identified as belonging to the genus *Vibrio*, we mapped these ASVs onto a reference phylogenetic tree of *Vibrio* spp. to reveal phylogenetic relationships. The three VDF-specific ASVs were not closely related to each other and were assigned to distinct clades, termed Vv1-Vv3 (Supplementary Fig. 4). Since these phylogenetic positions of these ASVs were determined based on the results of mapping, we further surveyed for their closely related species based on nucleotide sequence identities. ASV_3968 of Vv2 and ASV_4145 of Vv3 had identical nucleotide sequences to 16S rRNA genes from *Vibrio penaecida* and of *V. galatheae, V. hyugaensis, V. alginolyticus, V. harveyi,* and *V. nartriegens* (Table 1). Specifically, the Vv2 clade contained *V. penaecida*, while Vv3 clade included *V. alginolyticus*, *V. harveyi*, and *V. natriegens* (Supplementary Fig. 4).

**Table 1.** Phylogenetic and taxonomic assignment of VDF-specific and -nonspecific ASVs. Regarding ASVs whose sequence showed 100% identity with cultured bacteria deposited in RefSeq NR database, the hit bacterium is indicated with the corresponding accession number.

(A) *Vibrio* ASVs
| Clade | ASV_ID | The closest isolates | Accession ID of the closest isolates |
| --- | --- | --- | --- |
| Nv1 | ASV_516 | *Vibrio gigantis* | NR_114910 |
|  | ASV_694 |  |  |
|  | ASV_1745 |  |  |
| Nv2 | ASV_208 |  |  |
|  | ASV_3278 |  |  |
|  | ASV_3753 | *Vibrio chagasii* | NR_117891 |
| Vv1 | ASV_3626 |  |  |
| Vv2 | ASV_3968 | *Vibrio penaeicida* | NR_042121, NR_113790 |
| Vv3 | ASV_4145 | *Vibrio galathea* | NR_147758 |
|  | ASV_4145 | *Vibrio hyugaensis* | NR_145569 |
|  | ASV_4145 | *Vibrio alginolyticus* | NR_044825, NR_117895,<br>NR_122060 |
|  | ASV_4145 | *Vibrio harveyi* | NR_043165, NR_113784 |
|  | ASV_4145 | *Vibrio natriegens* | NR_026124, NR_113786,<br>NR_117890 |

(B) *Pseudoalteromonas* ASVs
| Clade | ASV_ID | The closest isolates | Accession ID of the closest isolates |
| --- | --- | --- | --- |
| Na1 | ASV_927 | *Pseudoalteromonas phenolica* | NR_113299 |
| Na2 | ASV_335 | *Pseudoalteromonas marina* | NR_042981 |
| Va | ASV_1731 |  |  |
|  | ASV_4400 |  |  |

The 5 VDF-nonspecific abundant *Vibrio* ASVs were grouped into two clades, termed Nv1 and Nv2 (Supplementary Fig. 4). Nv1 clade included ASV_516, ASV_694, and ASV_1745, while Nv2 clade included ASV_208, ASV_3278, and ASV_3753 (Table 1). ASV_516 (Nv1) and ASV_3753 (Nv2) show 100% nucleotide sequence identity with *V. gigantis* and *V. chagasii*, respectively (Table 1), suggesting that multiple ASVs within Nv1 and Nv2 are intraspecies populations of *V. gigantis* and *V. chagasii*, respectively. Due to the absence of 16S rRNA gene sequences longer than 1500 bp for *V. gigantis* and *V. chagasii* in public databases, these species were not included in the reference phylogenetic tree (Supplementary Fig. 4).

The VDF-specific and VDF-nonspecific *Vibrio* spp. possess distinct ecological characteristics. *V. penaecida* (Vv2) and *V. harveyi* (Vv3) are known pathogens of tiger prawn (Ishimaru *et al*., 1995) and bony fish such as sea bream and amberjack (Austin and Zhang, 2006; Monno *et al*., 2018), respectively. *V. gigantis (*Nv1) is an opportunistic pathogen of European seabass (Yilmaz *et al*., 2023) and has been shown to respond to intracellular components of uninfected *H. akashiwo* cells in our previous study (Takebe *et al*., 2024). *V. chagasii* (Nv2) is a known pathogen of oysters and mussels (Romero *et al*., 2014; Liang *et al*., 2019). Although no identical nucleotide sequences to ASV_3626 (Vv1) are available in the database, its mapped position suggests a closer relation to *Vibrio profundi* TP187, originally isolated from a seamount in the tropical western Pacific (Zhang *et al*., 2019) (Supplementary Fig. 4). Similarly, the VDF-specific abundant Pseudoalteromonadaceae ASVs (ASV_1731 and ASV_4400; *Pseudoalteromonas*) clustered together in a single clade (Va), distinct from the positions of the VDF-nonspecific Pseudoalteromonadaceae ASVs in the reference phylogenetic tree (Supplementary Fig. 5). The VDF-specific ASVs are closely related to *Pseudoalteromonas aliena* or *Pseudoalteromonas arctica* (Supplementary Fig. 5), although no identical 16S rRNA gene sequences to these species are available. In contrast, two VDF-nonspecific ASVs, ASV_927 (Na1) and ASV_335 (Na2), were placed in separate clades. ASV_335 and ASV_927 showed 100% nucleotide sequence identity with 16S rRNA genes of *Pseudoalteromonas marina* and *P. phenolica*, respectively (Table 1). The Na1 clade included *P. phenolica*. Additionally, the sequence of *P. marina* (CP023558) clustered with CP019162 *Pseudoalteromonas* sp. with >99.7% sequence identity, and CP019162 was included in the Na2 cluster, indicating *P. marina* also belonged to Na2 (Supplementary Fig. 5). *P. marina* is a marine species known for forming biofilms that facilitate the settlement of *Mytilus coruscus* larvae (Peng *et al*., 2020) and responding to intracellular components of uninfected *H. akashiwo* cells (Takebe *et al*., 2024). *P. phenolica* is noted for producing an antibiotic against certain bacteria (Isnansetyo and Kamei, 2003).

For the *Psychrobirium* ASVs, only VDF-specific lineage (ASV_1418) was identified in our experiment, so mapping to a reference phylogenetic tree of closely related species was not performed. However, ASV_1418 was found to be identical to 16S rRNA gene sequences of *Psychrobium conchae*. Close relatives of this species found associated with sea bream eggs, although its pathogenicity has not been reported (Najafpour *et al*., 2022).

It is important to observe whether the association of these abundant ASVs with *H. akashiwo* derived compounds observed in microcosm experiments also occurs in natural environments. We investigated the dynamics of VDF-specific and VDF-nonspecific abundant ASVs in natural bloom samples containing *H. akashiwo* collected in Monterey Bay, USA (Nowinski *et al*., 2019). We found a significant positive correlation between the relative abundance of VDF-specific *Pseudoalteromonas* ASV_1731 (Va) and VDF-nonspecific *Pseudoalteromonas* ASV_335 (Na2) with that of *H. akashiwo* (Supplementary Table 3). *H. akashiwo* increased during October 26-31 and November 5-7, 2016 and both ASV_335 and ASV_1731 also increased on November 5, 2016 (Supplementary Fig. 6). Although VDF-specific *Vibrio* ASVs, ASV_3968 and ASV_4145, as well as VDF-specific *Psychrobirium* ASV_1418, were detected during the bloom, the correlation between their dynamics and *H. akashiwo* was not statistically significant (Supplementary Table 3).

### 3.6. Difference in metabolic capacity between Vv clade and Nv clade

To gain insight into the metabolic capacities underpinning the increase of VDF-specific populations, we performed a comparative genomics analysis based on the composition of KEGG modules (Muto *et al*., 2013). We excluded *Psychrobirium* and *Pseudoalteromonas* from further analyses, as no VDF-nonspecific ASVs were found for *Psychrobirium*, and only one genome was available for close relatives of VDF-specific and VDF-nonspecific ASVs within *Pseudoalteromonas*. Consequently, we focused on *Vibrio*-spp., which included both VDF-specific (Vv2 and Vv3 clades) and VDF-nonspecific (Nv1 and Nv2 clades) populations, with available genomes (5, 44, 2, and 2 genomes, respectively) (Supplementary Table 4). Note that no genome was available for the ASV of the Vv1 clade. A total of 248 KEGG modules were identified in at least one genome, with varying patterns of module presence among the four clades (Supplementary Table 5). In particular, five modules were frequently present (> 70%) in the Vv2 and Vv3 clades (46 genomes), but not detected in the Nv clades (Fig. 4), suggesting different metabolic capability between Vv clades and Nv clades. These modules included- (i) biosynthesis of ectoine (M00033), an osmolyte, (ii) Branched−chain amino acid transport system (M00237) for branched-chain amino acid acquisition, (iii) Phosphotransferase (PTS) system cellobiose−specific II component (M00275) for importing and phosphorylating cellobiose, (iv) Nitrate/nitrite transport system (M00438) for nitrogen source acquisition, and (v) CreC−CreB (phosphate regulation) two−component regulatory system (M00449) for environmental condition recognition, such as carbon supply and oxygen availability (Wanner and Wilmes-Riesenberg, 1992; Avison *et al*., 2001), and specific cellular activities, such as biofilm formation (Zamorano *et al*., 2014) and motility (Huang *et al*., 2017) (Fig. 4). The branched-chain amino acid transport system was exclusive to Vv2 and Vv3 clade genomes, while the valine/isoleucine biosynthesis module (M00019) was present in genomes across all the four clades, indicating that those branched-chain amino acids might be biosynthesized but not essential amino acids (Supplementary Table 5). Furthermore, over 97% of Vv3 clade genomes (n=44) encoded the leucine degradation module (M00036), which converts leucine into acetyl-CoA, potentially linking it to the branched-chain amino acid transport system (Supplementary Table 5). Some of those modules were also detected in genomes of other *Vibrio* populations corresponding to ASVs that were not abundant in any treatment (Supplementary Table 6).

**Fig. 4.**
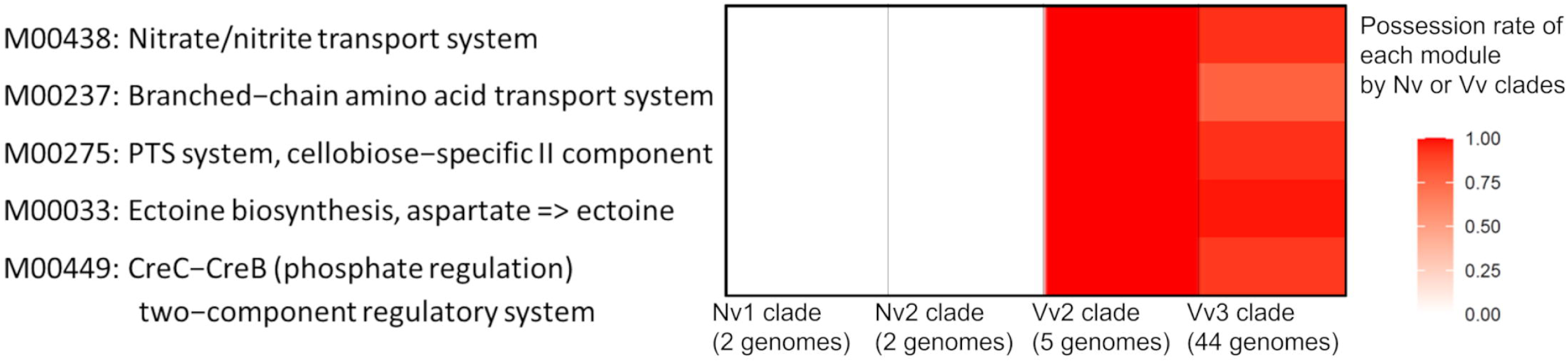
The distribution of KEGG modules in VDF-specific *Vibrio* (Vv) clades and VDF-nonspecific (Nv) clades. KEGG modules that are frequently (> 70%) detected in Vv clades genomes but not in Nv clades are shown. The heat map illustrates the possession rate of these KEGG modules across each clade.

### 3.7. Metabolites accumulated in VDF

We investigate whether the organic matter composition in VDF includes substrates specific to the metabolic capacities of the Vv clades using GC-MS analysis. Compared to the composition of low molecular weight (MW) compounds in f/2 medium, we detected 44 low MW compounds that were significantly more dominant in either of the three treatments (Supplementary Table 7). These compounds were likely derived from *H. akashiwo* cells rather than from the medium. The relative abundance of these 44 compounds varied among the different DFs (Supplementary Fig. 7). Especially, 15 of these compounds were at least twice as abundant in VDF as in the other DFs, suggesting a specific enrichment of these 15 compounds in the VDF (Fig. 5).

**Fig. 5.**
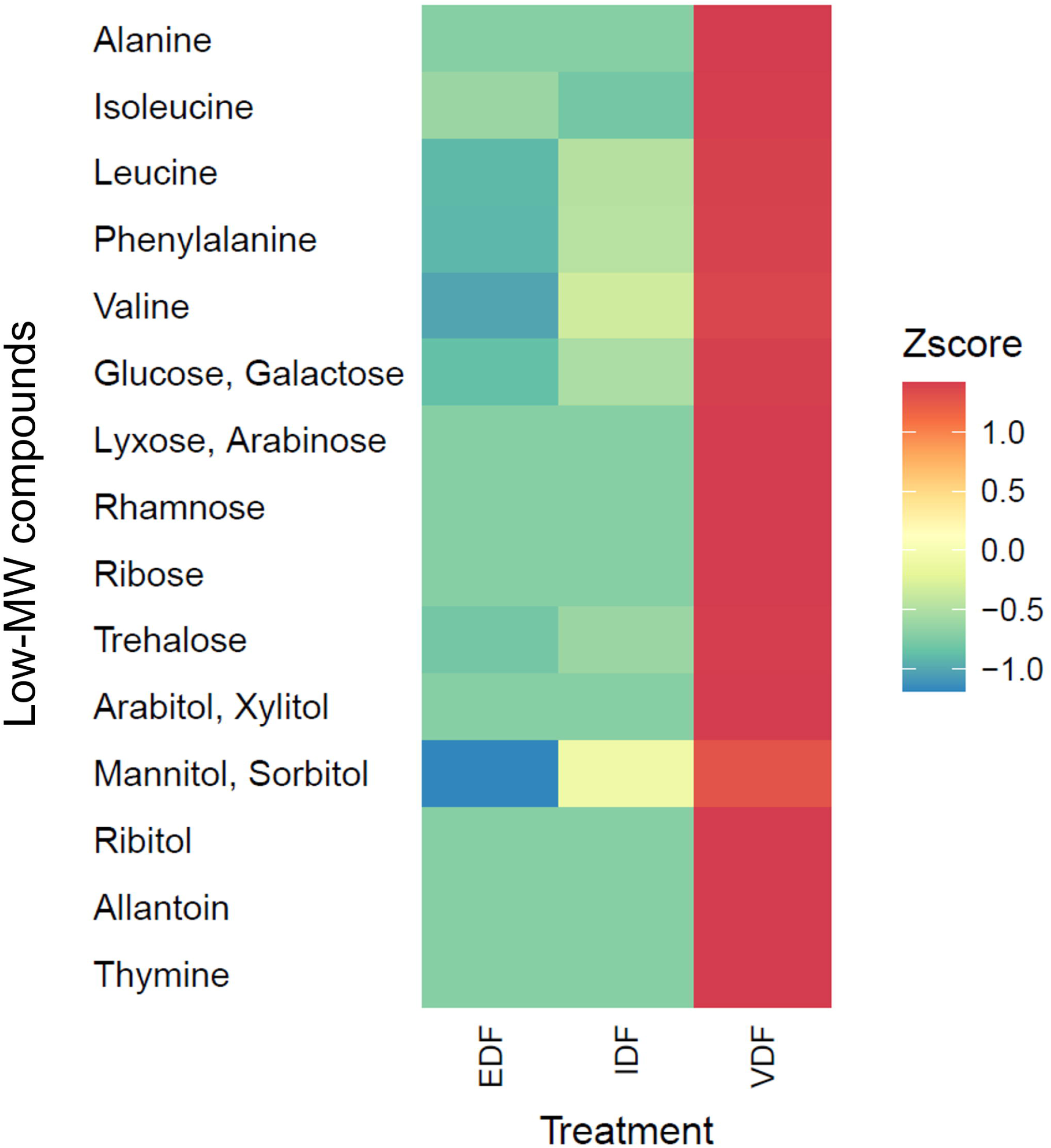
Comparison of relative abundance of low-molecular weight (MW) compounds enriched in VDF among VDF, IDF, and EDF. Only low-MW compounds that were at least twice as abundant in VDF than in IDF and EDF are presented. The relative abundance of each compound was calculated by dividing its peak area by that of internal control (2-isopropylmalate). The relative abundance was normalized with the carbon concentration of each sample enabling comparison between samples of different carbon concentration. The normalized relative abundance of each compound in each treatment was converted to z score.

The 15 low-MW compounds enriched in the VDF can be divided into four types; amino acids (alanine, isoleucine/allo-isoleucine, leucine, valine, and phenylalanine), sugars (glucose/galactose, ribose, lyxose/arabinose, rhamnose, and trehalose), sugar alcohols (mannitol/sorbitol, arabitol/xylitol, and ribitol), and nucleotide-derivatives (thymine and allantoin) (Fig. 5). Compounds separated by a slash represent those that could not be distinguished by GC-MS. Notably, all the branched-chain amino acids were specifically enriched in the VDF.

## 4. Discussion

In this study, we demonstrated that lysates released from *H. akashiwo* virocells promoted the growth of different prokaryotic populations compared to intracellular components or exudates from uninfected *H. akashiwo* cells (Figures 2 and 3). The virocell-specific prokaryotic populations included ASVs from the genera *Vibrio*, *Pseudoalteromonas*, and *Psychrobirium* (Supplementary Table 2). While our previous microcosm experiments showed that certain populations in these genera responded to uninfected *H. akashiwo* cells (Takebe *et al*., 2020, 2024), this study revealed that the preference for *H. akashiwo* virocells or uninfected cells varied even among ASVs within the same genera (Table 1). Interactions between *H. akashiwo* virocells and certain prokaryotic populations, and those between uninfected *H. akashiwo* cells and other prokaryotic populations, likely occur not only under laboratory conditions but also in the natural environments. This is supported by the observed correlations between the dynamics of VDF-specific and VDF-nonspecific *Pseudoalteromonas* populations and *H. akashiwo* in Monterey Bay, US (Supplementary Fig. 6). Although the correlation with *H. akashiwo* dynamics was not statistically significant for VDF-specific and VDF-nonspecific *Vibrio* populations, these groups were also present during the bloom. Given the detectable concentrations of HaV during *H. akashiwo* blooms in previous studies, it is likely that *H. akashiwo* cells in natural environments are exposed to HaV infection, leading to the lysis of at least some of the cells (Tarutani *et al*., 2000). Therefore, natural *H. akashiwo* blooms are likely a mixture of uninfected cells and virocells and increases in both VDF-specific and VDF-nonspecific populations in natural blooms would be rational. However, the proportion of *H. akashiwo* virocells in natural blooms remain unclear, as there is no plausible method for specific and quantitative detection. This remains a subject for future research.

Virocell-specific prokaryotic responses have been reported in several eukaryotic microalgae (Sheik *et al*., 2013) and cyanobacteria (Xiao *et al*., 2020). Experiments using ^13^C-labeled tracers or quantification of DOM in liquid media suggest that DOM from the virocells is utilized by specific, heterotrophic bacteria (Sheik *et al*., 2013). However, the specific compounds from microalgal virocells that promote the growth of specific prokaryotic communities remained unclear in these studies. Genomic analyses of *Vibrio* spp., which are closely related or phylogenetically similar to the VDF-specific *Vibrio* ASVs (ASV_3968 and ASV_4145), reveal that these strains specifically encode a branched-chain amino acid transport system (M00237) not detected in VDF-nonspecific *Vibrio* populations (Fig. 4). Additionally, these strains also encode a valine/isoleucine biosynthesis module (M00019) (Supplementary Table 5). The biosynthesis of valine and isoleucine begins with pyruvate, a key metabolite involved in the biosynthesis of fatty acids, lipids, and isoprenoids. Consequently, the biosynthesis of branched-chain amino acids might compete with the biosynthesis of other biologically essential compounds. Although acquisition of branched-chain amino acids from the environments consumes ATP-since the transporter belongs to the ATP binding cassette family (Matsubara *et al*., 1992), this system might provide an additional source of amino acids, thereby conserving pyruvate in VDF-specific *Vibrio* populations. Furthermore, all genomes of Vv3 clade, except one, encode leucine degradation modules (M00036) (Supplementary Table 5). This pathway degrades one of the branched-chain amino acids into acetyl-CoA, which is also a key substrate used for various metabolic pathways, including fatty acid biosynthesis and respiration. Therefore, acquiring branched-chain amino acids from VDF could support the synthesis of various compounds or energy conservation in the Vv3 clade, promoting its growth. Notably, the relative abundances of branched-chain amino acids, such as leucine, isoleucine, and valine were higher in the VDF than in the IDF and EDF (Fig. 5). These VDF-specific compounds may serve as substrates for the branched chain amino acid transport system in the *Vibrio* populations. This is the first report to identify candidate metabolic compounds from virocells that might affect the surrounding prokaryotic community structure. It is important to recognize that the abundance of prokaryotic population is determined by complex interplay off factors, including both nutrients and environmental conditions. Thus, the increase in specific populations cannot be attributed to a single factor alone. While *H. akashiwo* virocell-derived VDF may regulate specific *Vibrio* populations, it does not necessarily promote the increase of all *Vibrio* strains with the branched-chain amino acid transport system in the present microcosm experiment (Supplementary Table 6). It remains possible that the less abundant populations could become more prevalent abundant under different conditions with VDF-derived compounds. Other metabolic features in prokaryotic populations might also influence their dynamics in the presence of *H. akashiwo* virocell-derived compounds. In addition to branched-chain amino acids transport system, four additional metabolic modules were prevalent in the Vv clades: ectoine biosynthesis, phosphotransferase system, cellobiose−specific II component, nitrate/nitrite transport system, and creC−creB two−component regulatory system (Fig. 4). Especially, creC, a sensory kinase in the creC-creB regulatory system, is known to respond changes in carbon supply (Wanner and Wilmes-Riesenberg, 1992; Avison *et al*., 2001). Sugars such as glucose/galactose, ribose, lyxose/arabinose, rhamnose, and trehalose, analyzed by GC-MS were relatively enriched in *H. akashiwo* VDF when compared to other DFs. Changes in the composition of carbon sources due to *H. akashiwo* virocell lysis might be sensed by creC kinase, potentially triggering a response from the creB regulatory factor, affecting activities of VDF-specific *Vibrio* populations, such as biofilm formation and motility (Avison *et al*., 2001; Huang *et al*., 2017). Currently, we have not identified plausible links of virocell-derived compounds and the modules for ectoine biosynthesis, phosphotransferase system, cellobiose−specific II component, and nitrate/nitrite transport system. Further research is needed to elucidate these potential connections.

The effects of some low-MW compounds enriched in the virocell lysates on viral particle production remains unclear. Viral particle production requires nucleotides and amino acids for DNA/RNA and protein synthesis, respectively. While it has been reported that the major capsid protein of *Chlorella* viruses, which infect certain strains of the green alga *Chlorella*, is modified by glycans (Van Etten *et al*., 2017). However, no studies have linked sugar alcohols to the production of microalgal viruses. Amino acids, sugars, sugar alcohols, and nucleotide derivatives are likely biosynthesized by some algal cells, as they have been detected in cellular compounds, mucilage sheaths, and/or excreted compounds from various algal cells (Maruo *et al*., 1965; Reed *et al*., 1985; Karsten *et al*., 1990; Watanabe *et al*., 2006; Kosugi *et al*., 2013; Groisillier *et al*., 2014; Rosenwasser *et al*., 2014; Hirth *et al*., 2017; Hano and Tomaru, 2023). However, lyxose and allantoin are exceptions. Allantoin, an intermediate metabolite in purine catabolism and common in plants (Kaur *et al*., 2021), has not been reported in algae. To confirm this, we searched for homologous proteins involved in *Arabidopsis* allantoin biosynthesis from xanthin (Accession nos. AT4G34890, AT4G34900, AT2G26230, and AT5G58220) in the transcriptome data of uninfected *H. akashiwo* CCMP452 (https://dx.doi.org/10.6084/m9.figshare.3840153.v3), but found no homologues. Similarly, homologous proteins were not found in the genome of HaV01 (https://www.ncbi.nlm.nih.gov/datasets/taxonomy/97195/). Whether the virocell-specific low-MW compound identified as allantoin is indeed genuine allantoin or an as-yet-unknown compound that is difficulty to be distinguished from allantoin remains to be confirmed. Future studies should clarify whether it is true allantoin and elucidate its role, if any in viral particle production within *H. akashiwo* virocells.

Finally, the virocell-specific *Vibrio* populations or their close relatives are pathogens of tiger prawns and bony fish including sea bream and amberjack, while virocell-nonspecific ones infect oyster and mussel (Table 1). Therefore, changes in prokaryotic dynamics caused by viral infections in *H. aksahiwo* might lead to negative impacts in marine environments. Actually, blooms of *H. akashiwo* are known to damage sea bream, amberjack, and oysters (Keppler *et al*., 2005; Mehdizadeh Allaf, 2023). Proposed mechanisms for these impacts include excessive mucous secretion, and production of reactive oxygen species, organic toxin, and hemolytic compounds from *H. akashiwo* cells during blooms; however, these mechanisms remain to be elucidated (Mehdizadeh Allaf, 2023). Although not reported previously, these negative impacts might be indirectly mediated by *H. akashiwo*; possibility through bacteria that respond the alga. The biological and biogeochemical links between viral infections in *H. akashiwo* and the marine material cycling and dynamics of higher trophic-level consumers, such as fish and crustaceans could be significant.

## 5. Conclusion

In the present study, we investigated effects of lysates from *H. akashiwo* virocells on the dynamics of coastal prokaryotic populations. Our data suggested that the virocell-specific prokaryotes had certain bacterial populations that appear to utilize specific organic matters enriched only in virocells. indicating that changes in organic compounds induced by viral infection promote growth of prokaryotes with specific metabolic capacities. Additionally, as the population included pathogenic bacteria of fish or crustacean, viral infection to *H. akashiwo* can indirectly affect the dynamics of higher-trophic level consumers. Considering that *H. akashiwo* is a widely distributed bloom-forming species and viral infection is a crucial factor in terminating their blooms, understanding in these dynamics is crucial for assessing the broader ecological impacts of *H. akashiwo* blooms and their associated viral infections.

## Acknowledgements

Computational analyses were partly performed at the Super Computer System, Institute for Chemical Research, Kyoto University. This study was supported by Grants-in-Aid for Scientific Research (No. 21H05057, and No. 21J14854) from the Japan Society for the Promotion of Science (JSPS). CRediT authorship contribution statement Hiroaki Takebe: Funding acquisition, conceptualization, investigation, formal analysis, visualization, and writing (original draft). Haruna Hiromoto: Investigation. Tetsuhiro Watanabe: Resources. Keigo Yamamoto: Resources. Keizo Nagasaki: Resources. Ryoma Kamikawa: Conceptualization, supervision, and writing (review and editing). Takashi Yoshida: Funding acquisition, project administration, conceptualization, supervision, and writing (review and editing).

## Data availability

The sample information obtained in this study was deposited in the DNA Data Bank of Japan (DDBJ) under project number PRJDB18787. Raw sequence reads can be found under accession number DRA019308.

