## Supplementary Figures for "Viral infection to the raphidophycean alga *Heterosigma akashiwo* affects both intracellular organic matter composition and dynamics of a coastal prokaryotic community"

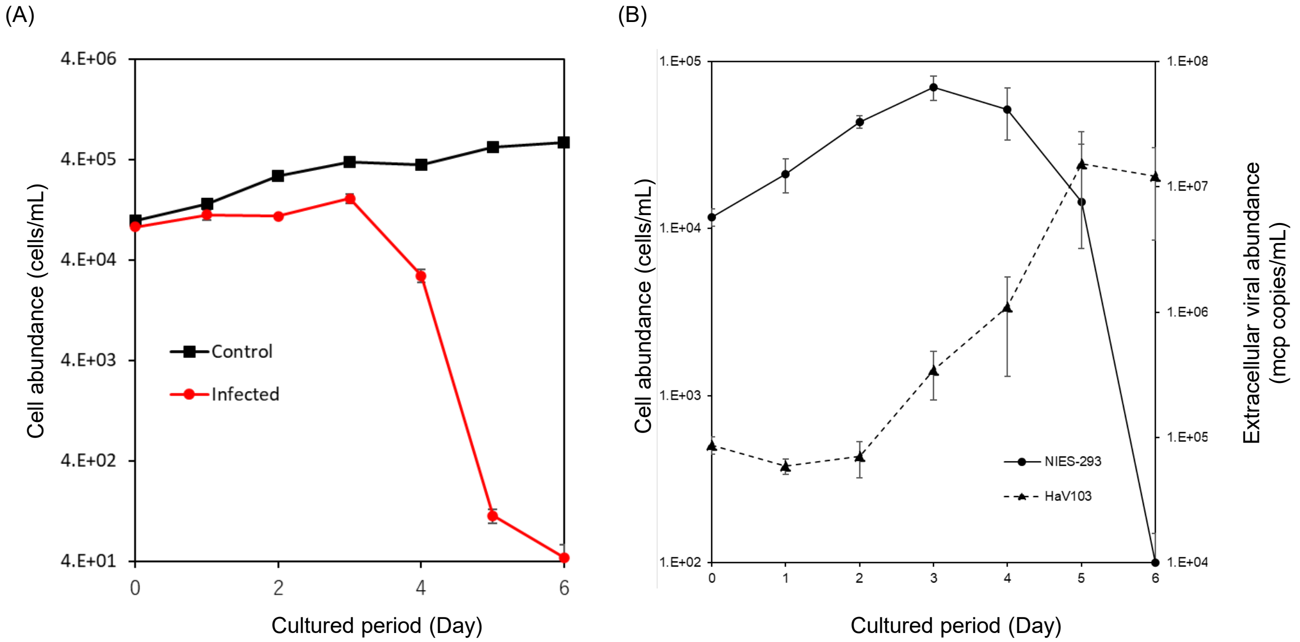


**Supplementary Fig. 1.** Shifts in abundance of *H. akashiwo* NIES-293 and *H. akashiwo* virus HaV103 during infection experiments. (A) uninfected control and infected culture. (B) 10-times diluted infected culture constructed for monitoring HaV103 abundance. Cell counts were obtained using flow cytometry and HaV103 abundance was analyzed by quantitative PCR targeting major capsid gene (mcp). (A) The average cell abundance in the triplicate measurements is shown. (B) The average cell and viral abundance in the triplicate flasks are shown. Error bars indicate standard deviation.

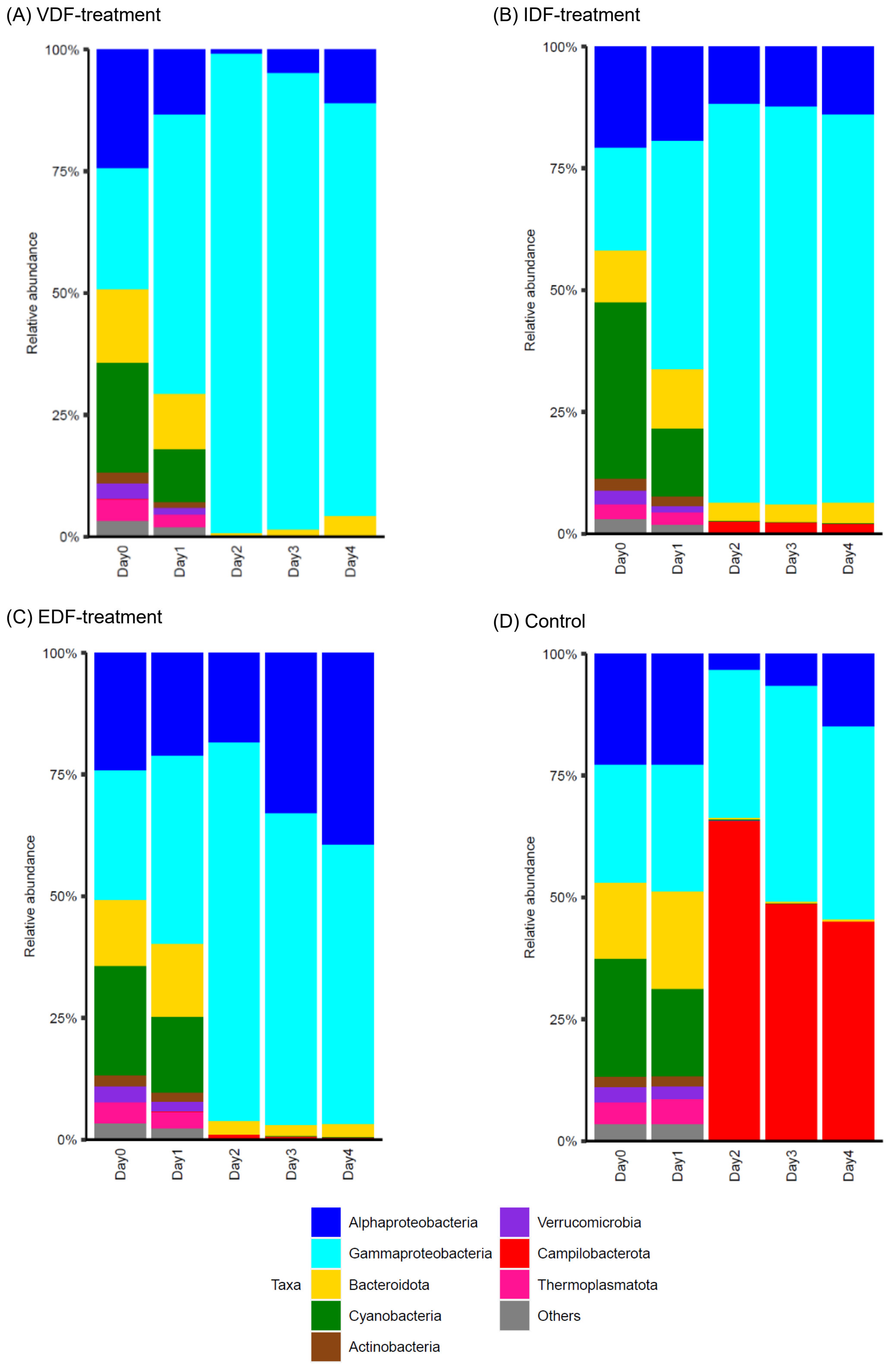


**Supplementary Fig. 2.** Relative abundance of phylum-level (class-level for proteobacteria) phylogenetic groups in the microcosm samples. For each treatment, averaged relative abundance in the triplicate flasks is shown.

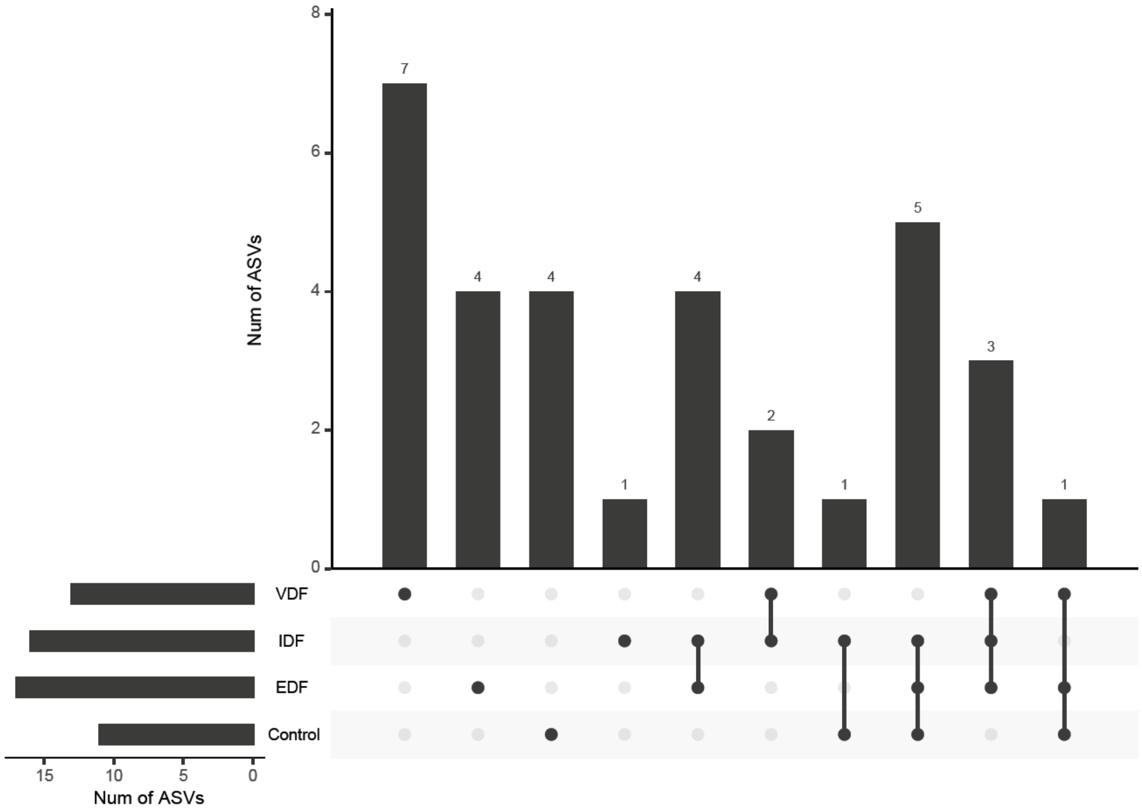


**Supplementary Fig. 3**. UpSet plot indicating the distribution pattern of all the abundant ASVs among the treatments. The number shown above bar graph indicates those of abundant ASVs detected in each treatment.

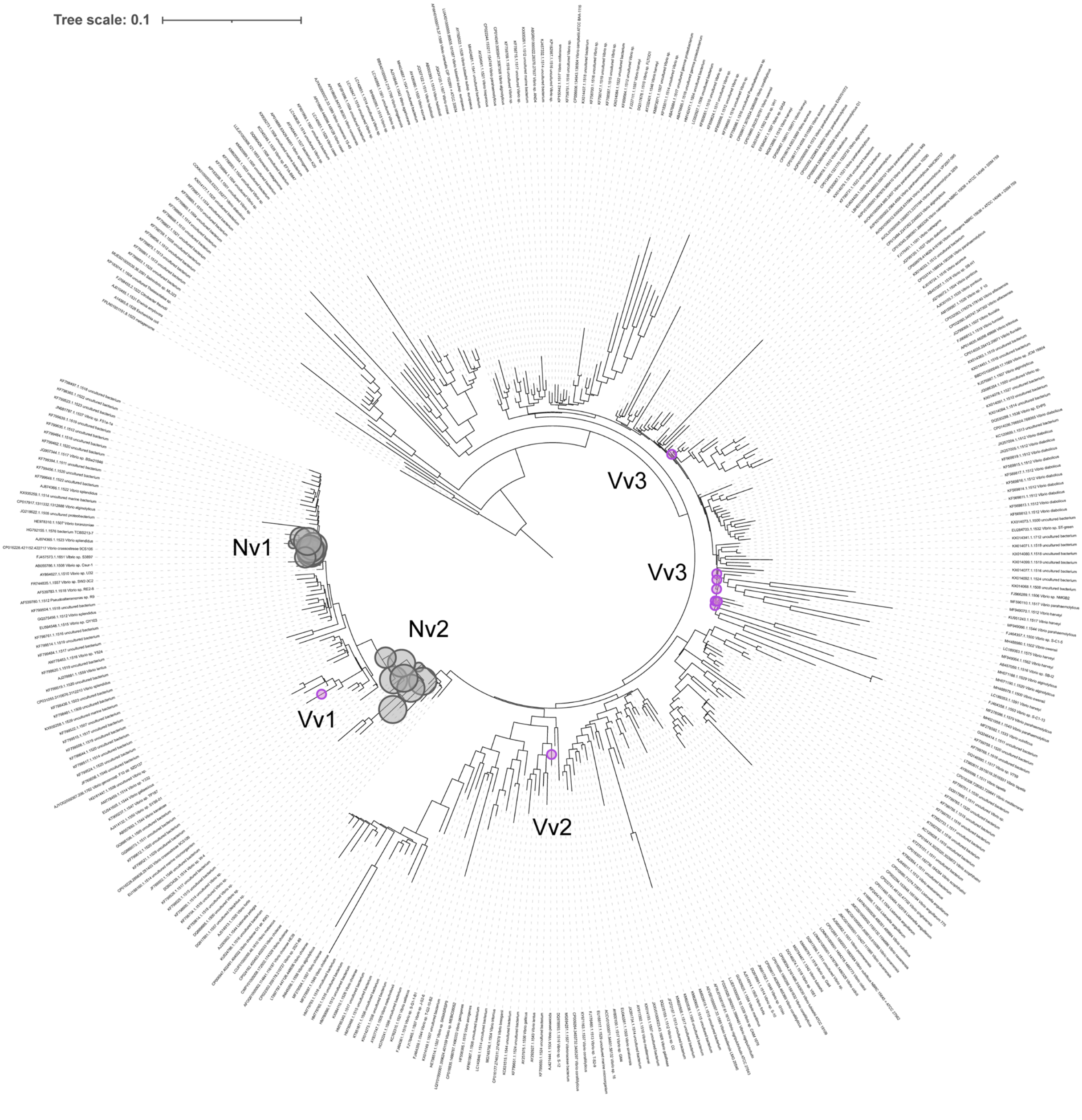


**Supplementary Fig. 4.** Phylogenetic placement of Vibrionaceae abundant ASVs on reference tree constructed with nearly complete 16S rRNA genes. The reference phylogenetic trees were constructed using the approximately-maximum likelihood method. The purple circles indicate the nodes where VDF-specific ASVs were mapped and the gray circles did the nodes where VDF-nonspecific ASVs were mapped. Vv1; ASV_3626. Vv2; ASV_3968. Vv3; ASV_4145. Nv1; ASV_516, ASV_694, and ASV_1745. Nv2; ASV_208, ASV_3278, and ASV_1745.

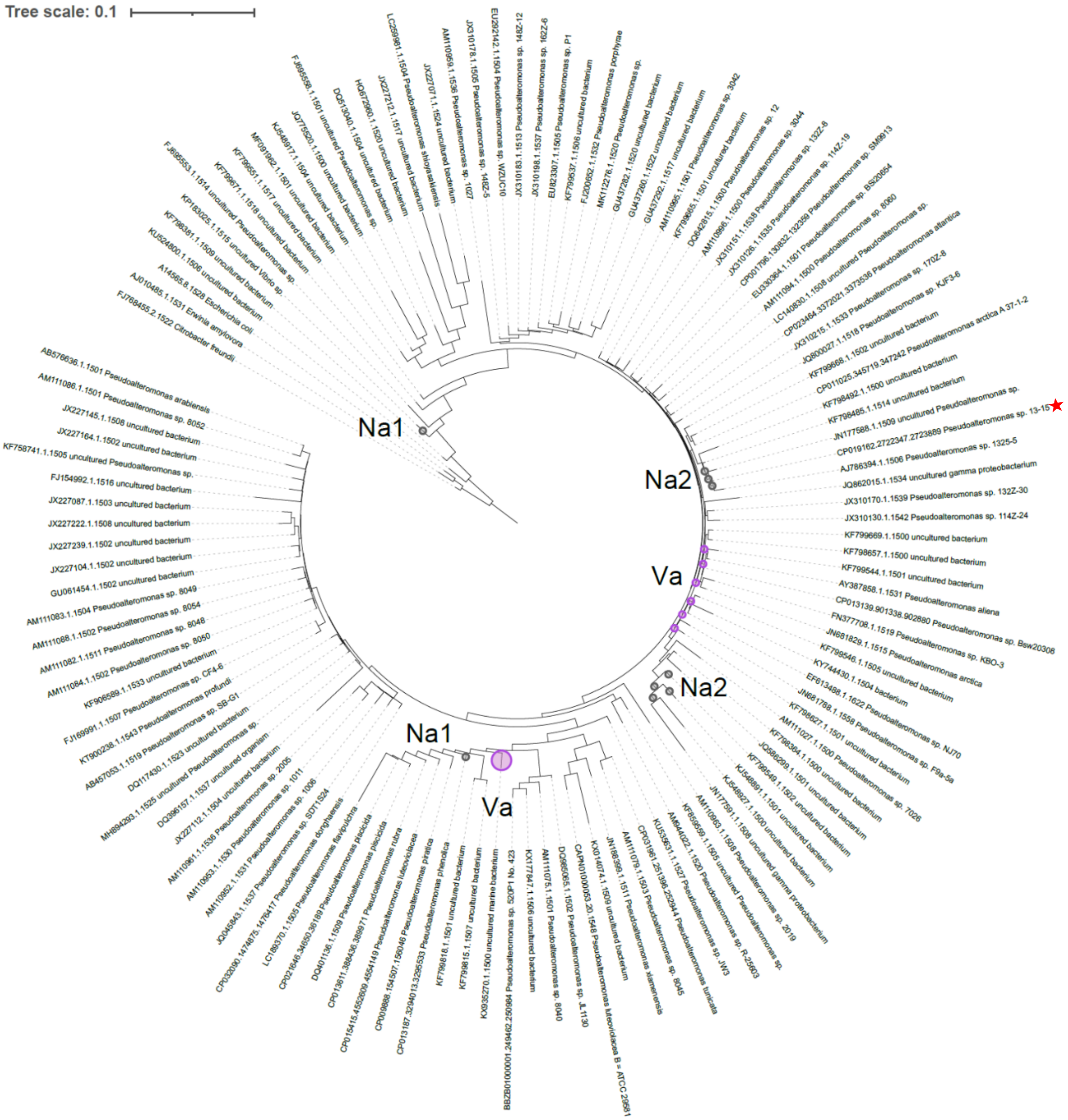


**Supplementary Fig. 5.** Phylogenetic placement of Pseudoalteromonadaceae abundant ASVs on reference tree constructed with nearly complete 16S rRNA genes. The reference phylogenetic trees were constructed using the approximately-maximum likelihood method. The purple circles indicate the nodes where VDF-specific ASVs were mapped and the gray circles did the nodes where VDF-nonspecific ASVs were mapped. Va; ASV_1731 and ASV_4400. Na1; ASV_335. Na2; ASV_927. The sequence CP019162.2722347.2723889_*Pseudoalteromonas*_sp._13-15, indicated by the red star, clustered with the 16S rRNA gene sequence of *P. marina* (CP023558) with >99.7% sequence identity.

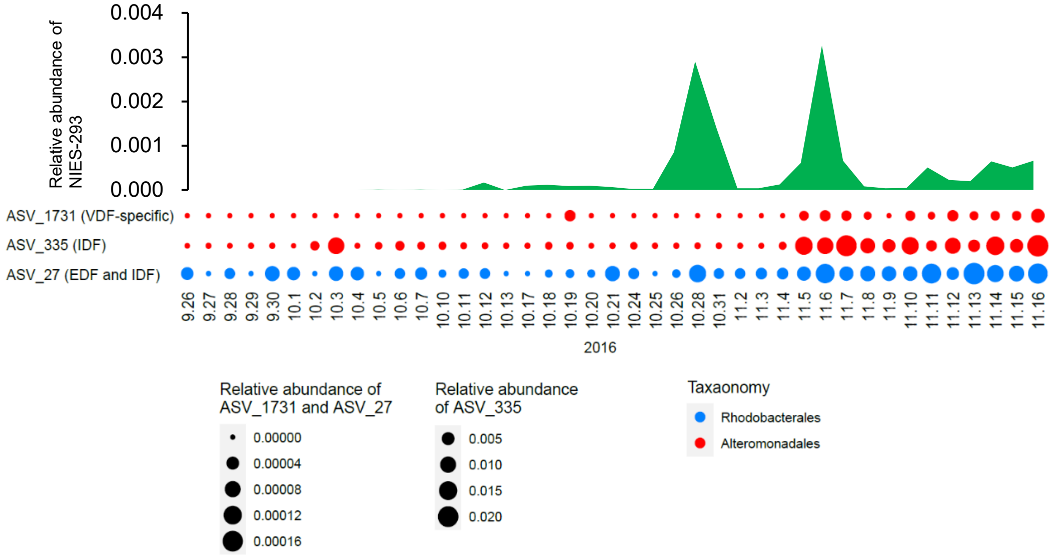


**Supplementary Fig. 6.** Co-occurrence dynamics of the *H. akashiwo* and the abundant ASVs during the microcosm experiments in the natural bloom samples. The microbiome sequence datasets almost daily collected between September 26–November 16, 2016 (Nowinski *et al.,* 2019) were used. Relative abundance of *H. akashiwo* NIES-293 close relatives was calculated by mapping quality-controlled reads of 18S rRNA genes to the NIES-293 sequence with 100% identity using VSEARCH. Relative abundance of each ASV was calculated by mapping quality-controlled reads of 16S rRNA genes to the ASV sequence with 100% identity using VSEARCH. Co-occurrence dynamics are shown only if significantly positive correlation is detected (Spearman correlation; $r > 0.6$, $p < 0.01$ and $Q < 0.05$).

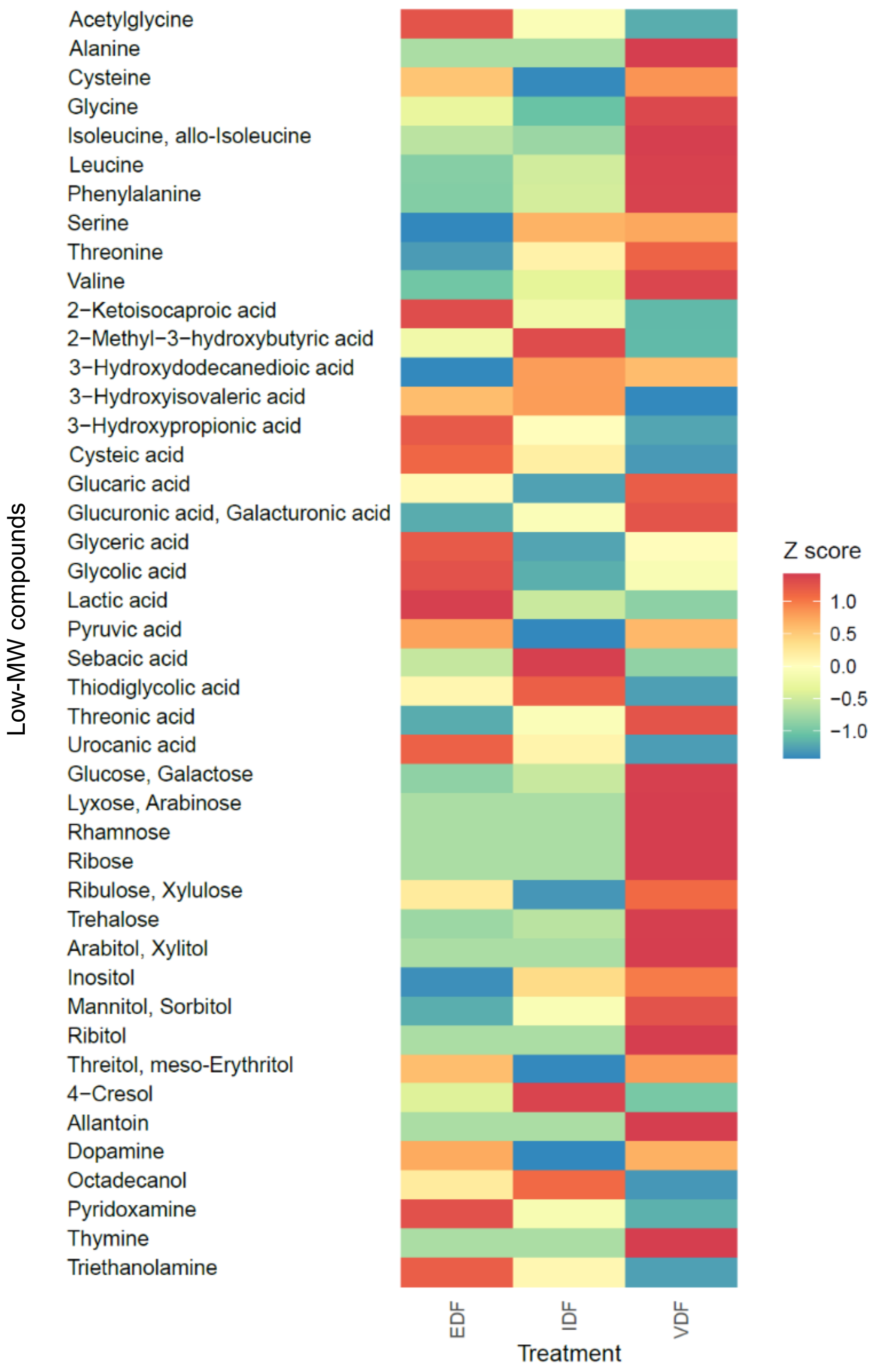


**Supplementary Fig. 7.** Comparison of the relative abundance of low-molecular weight (MW) compounds detected in VDF, IDF, and EDF by GC-MS analysis. Only low-MW compounds which showed higher relative abundance in any of VDF, IDF, or EDF than in control are presented. The relative abundance of each low-MW compound was dividing the peak area of the compound by that of internal control (2-isopropylmalate). The relative abundance was normalized with the carbon concentration of each sample enabling comparison between samples of different carbon concentration. The normalized relative abundance of each compound in each treatment was converted to z score.
